# Discovery of non-opioid receptor protein targets of fentanyl and remifentanil by affinity-based protein profiling in diverse animal model and human tissues

**DOI:** 10.1101/2025.02.20.634605

**Authors:** Vivian S. Lin, Kiall F. Suazo, Doo Nam Kim, Damon T. Leach, Agne Sveistyte, Jennifer Walker, Leo J. Gorham, Katherine J. Schultz, Kai-For Mo, Stephen J. Callister, Kelly G. Stratton, Gerard X. Lomas, William C. Nelson, Heather A. Colburn, Vanessa L. Paurus, Priscila M. Lalli, Ronald J. Moore, Samantha M. Powell, Oscar Rodriguez, John R. Cort, Aaron T. Wright

## Abstract

Synthetic opioids such as fentanyl and related analogs have been widely used for pain management. However, their negative side effects, including respiratory depression and high potential for addiction, underscore the need for a deeper understanding of fentanyl’s interactions with proteins throughout the human body. Fentanyl analogs bind and activate opioid receptors in the central and peripheral nervous systems, triggering numerous downstream signaling pathways. Increasingly, fentanyl has been shown to interact with non-opioid receptors, and elucidation of these non-canonical fentanyl-protein interactions may provide insights into the mechanisms contributing to fentanyl’s adverse effects and illuminate novel countermeasure strategies. To identify proteins in mammalian tissues that may interact with fentanyl, we designed and synthesized three affinity-based probes (AfBPs) that include the fentanyl core and feature a diazirine photoaffinity group and alkyne handle for click chemistry at different positions. Molecular docking simulations predicted that these AfBPs bind the mu opioid receptor similarly to fentanyl.

Affinity-based protein profiling using the FA-T1 probe *in vitro* in tissues from six animal species identified histamine N-methyltransferase (HNMT), endophilin-B1 (SH3GLB1), fructosamine-3-kinase (FN3K), cutA divalent cation tolerance analog (CUTA), and monoamine oxidase B (MAOB) among the top proteins that bind fentanyl in multiple species and tissue types. Molecular docking of fentanyl and remifentanil with these protein structures identified putative binding sites. The interaction of fentanyl with specific proteins was empirically assessed through protein structural analyses. These findings highlight potential fentanyl-protein interactions that may contribute to the acute and long-term impacts of fentanyl exposures.

## Introduction

The development of ultrapotent synthetic opioids in the mid-20^th^ century led to widespread usage of these drugs for anesthesia, sedation, and pain management. Fentanyl, a phenylpiperidine opioid narcotic, was first synthesized in 1960 and displays 50 to 100 times higher potency than morphine.^1^ Like all opioids, fentanyl exerts its physiological effects primarily through agonism of opioid receptors throughout the body, particularly the mu opioid receptor (MOR), with comparatively low affinity for the delta and kappa opioid receptors.^2^ Despite their effectiveness for sedation and analgesia, fentanyl and its analogs display serious negative side effects such as respiratory depression, brachycardia, and high addiction potential. Fatal overdose due to fentanyl is frequently caused by rapid onset of respiratory depression,^3-5^ with over 84,000 deaths in the United States attributed to fentanyl in 2022.^6^ Fentanyl’s ease of synthesis has also led to the widespread emergence of designer fentanyls; to date, more than 100 known fentanyl analogs have been identified, some of which display higher potencies than fentanyl itself.^7^ Naloxone, a MOR antagonist, can be used to reverse symptoms of opioid overdose. Increased availability of naloxone and community education has reduced fatal opioid overdose in recent years,^8^ although additional efforts are underway to improve upon this antidote and develop alternate countermeasures.^9^

As countermeasure development for acute opioid-induced respiratory depression advances, questions remain regarding effects of fentanyl that do not involve the opioid receptors. Fentanyl has been reported to bind human ether-a-go-go related gene (hERG),^10, 11^ serotonin (5-HT) receptors,^12, 13^ and dopamine receptor,^14^ muscarinic receptor,^15-17^ and adrenergic receptor subtypes.^14, 18-21^ Besides respiratory depression, wooden chest syndrome (WCS), characterized by chest muscle wall and diaphragm rigidity,^22^ and vocal cord closure (VCC)^23-25^ are life-threatening physiological phenomena observed in both humans and animal models of fentanyl overdose. WCS and VCC are generally unresponsive to naloxone treatment, suggesting they are mediated by mechanisms distinct from MOR agonism. Furthermore, the long-term health effects of high dose fentanyl exposures in overdose survivors remain virtually unknown. Increasingly, fentanyl’s impacts on organs beyond the brain are also being recognized, particularly for chronic opioid use.^26-28^ The potential involvement of non-opioid receptors in both acute and long-term adverse health impacts of ultrapotent synthetic opioids remains relatively understudied and merits further investigation. Strategies to identify protein receptors that interact with fentanyl and trigger signaling pathways contributing to adverse effects of fentanyl overdose will be important steps toward developing more effective treatments to combat the opioid crisis.

Affinity-based protein profiling (AfBPP) is a chemoproteomic technique that uses affinity-based probes (AfBPs) to covalently label receptors that interact with a ligand of interest by mimicking the ligand’s chemical structure or properties.^29^ This approach is particularly useful for profiling proteins of interest in complex mixtures due to its ability to enrich selected proteins from the proteome, providing more rapid and unbiased identification of proteins that may interact with the ligand in specific biological systems. AfBPs may contain photocrosslinking groups, such as a diazirine or benzophenone, to irreversibly and covalently attach the probe to proteins. A wide variety of AfBPs have previously been developed for protein-drug interaction discovery, including photoaffinity probes for propofol,^30^ methamphetamine,^31^ and morphine.^32^ However, despite the utility of the AfBPP approach for identifying novel protein-drug interactions and the need to better understand fentanyl-protein interactions beyond opioid receptor-mediated signaling, no AfBPs based on the structure of fentanyl have been reported to date.

To enable the discovery of protein-fentanyl interactions in complex proteomes, we designed three AfBPs for chemoproteomic profiling of fentanyl-binding proteins in tissue homogenates (**Figure 1A**). We envisioned that fentanyl-based AfBPs could be readily produced by incorporating a photoactivatable crosslinking group into different locations on the fentanyl core structure.^33^ The photoactivatable diazirine crosslinker forms a highly reactive carbene upon photolysis with ultraviolet (UV) light; this carbene will undergo an insertion reaction with nearby molecules, enabling the probe to be covalently and irreversibly attached to proteins to which it is bound.^34^ To enable flexible downstream conjugation of reporters to the probe-protein adduct, we also incorporated an alkyne “click” group for copper catalyzed azide-alkyne cycloaddition to attach either a fluorophore for polyacrylamide gel electrophoresis (SDS-PAGE) or biotin for biotin-avidin enrichment and mass spectrometry-based proteomic analysis.

**Figure 1.**
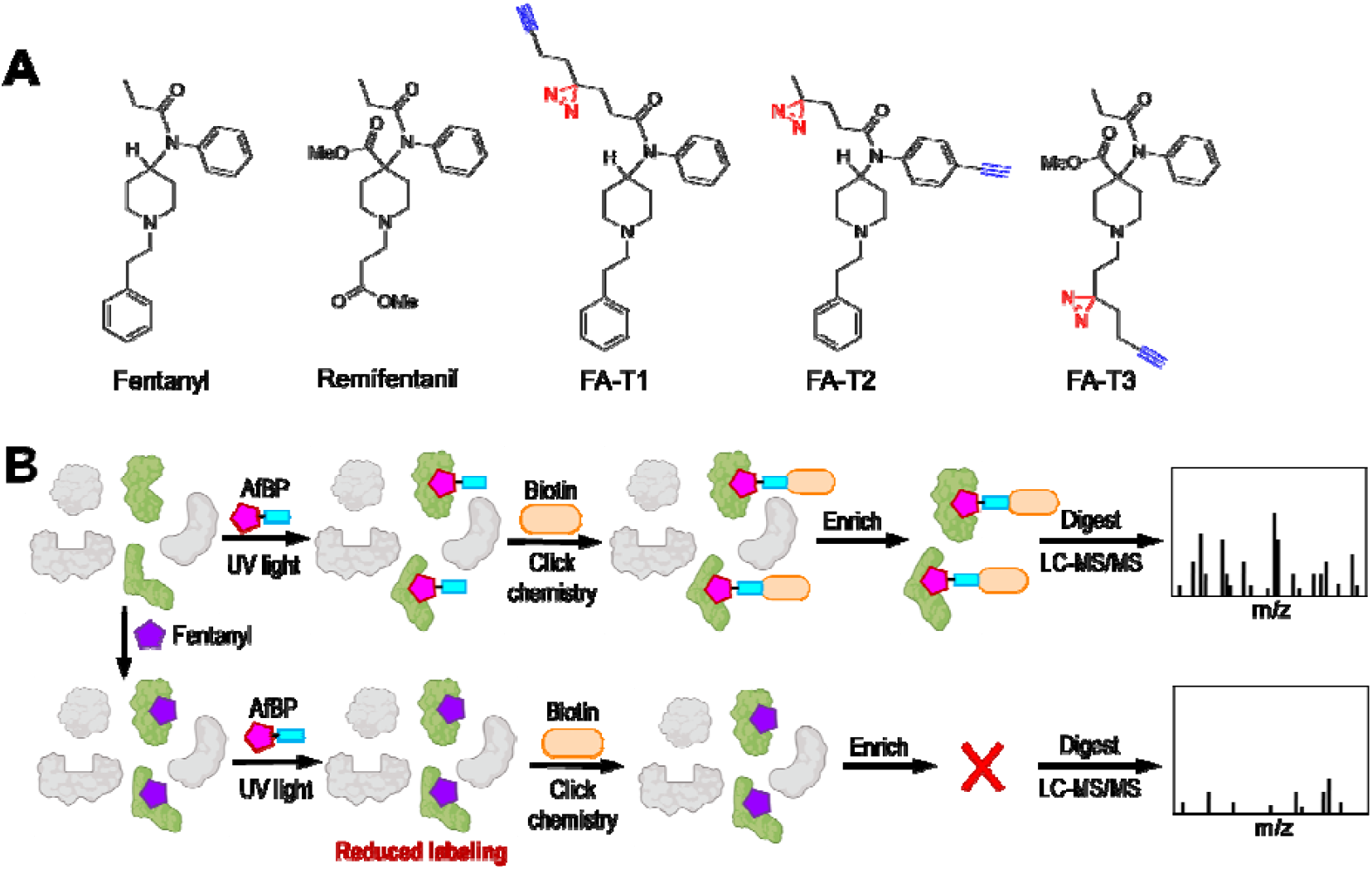
(A) Structures of fentanyl analogs and fentanyl affinity-based probe (AfBP) designs. (B) Competitive affinity-based protein profiling workflow for fentanyl-binding protein discovery. Fentanyl preincubation leads to reduced AfBP labeling of protein targets by occupying available binding sites.

To ensure that the AfBPP approach identifies proteins that interact with authentic fentanyl analogs, we pretreated the tissue homogenates with vehicle or fentanyl and remifentanil before labeling with the probes. This competitive AfBPP strategy allows for the fentanyl analogs to occupy binding sites in the proteome, blocking probe binding; high confidence protein targets are therefore proteins that show decreased AfBP labeling with competitor treatment (**Figure 1B**).^35, 36^ To advance our understanding of fentanyl’s impacts on the body, we profiled brain, heart, kidney, lung, and liver to identify potential fentanyl-binding proteins in these organs that may be affected during overdose. Furthermore, to better understand both shared and species-specific protein targets of fentanyl analogs in mammalian models, we profiled tissues from mouse, rat, guinea pig, ferret, and rhesus macaque, in addition to humans. A major challenge in toxicology is translating protein target relevance from animal models to humans. To meet this need, we aimed to generate the first comprehensive multi-organ, multi-species profile of fentanyl-interacting proteins to begin connecting potential molecular level interactions to observed physiological phenomena. A deeper understanding of fentanyl’s protein interactome beyond opioid receptors will provide an important first step toward clarifying the potential mechanisms by which ultrapotent synthetic opioids modulate signaling pathways and may lead to new strategies to address the adverse health impacts associated with opioid abuse and overdose.

## General materials and methods

### Computational methods

Molecular docking simulations were run using AutoDock Vina^37^ with a co-crystallized 3D structure of human MOR obtained from www.rcsb.org (PDB ID 5C1M). The center of mass coordinate for BU72, the co-crystallized morphinan agonist, was identified and subsequently used to define the center point of the docking search volume. BU72 was deleted prior to running the simulations. The probes were loaded as 3D structures in arbitrary poses, with docking simulations run for each. The top eight poses and corresponding binding scores were collected for fentanyl and each probe. Decoys were generated using DUD-E^38^ to serve as presumed negative binders of MOR for binding affinity score comparison.

To further assess experimental AfBP binding results, we performed docking simulations for fentanyl and remifentanil binding into muscarinic acetylcholine receptor M1 (CHRM1), histamine N-methyl transferase (HNMT), and monoamine oxidase B (MAOB). For ligand structure preparation, we optimized their geometries using Avogadro 1.2.0.^39^ Following the optimization, 50 standard global ensemble conformers were generated for each refined ligand structure using BCL.^40^ While AutoDock inherently evaluates multiple conformers of the provided ligand, our additional conformer sampling aimed to explore an even broader range of the ligand’s structural landscapes. Receptor structures were downloaded from www.rcsb.org (PDB ID 6ZFZ, 2AOT) and pre-processed using the ‘Dock Prep’ tool in UCSF ChimeraX.^41^ To enhance computational efficiency, receptor structures were kept rigid during the docking process, consistent with our previously established method.^42^ Unbiased “global” docking was ensured by adjusting grid limits to encompass the entire receptor. After running the AutoDock docking simulation, the top pose—defined by the lowest binding energy—was selected for further evaluation. The visualization of docked results and protein-ligand interaction analysis were carried out with ChimeraX^41^ and ProteinsPlus.^43^

### Probe synthesis

Synthesis, storage, and experimental use of controlled substances including Schedule I-II fentanyl analogs was carried out in accordance with Drug Enforcement Agency (DEA) and Washington State Department of Health (WDOH) regulations. Detailed synthetic methods are described in the Supporting Information.

### AfBPP sample preparation

Tissue homogenization and qualitative analysis of AfBP labeling in tissue homogenates by SDS-PAGE are described in the Supporting Information. For AfBPP proteomics sample preparation, proteome samples were removed from −70 °C storage and thawed on ice. Lysates were normalized to a final concentration of 2 mg/mL with cold 1 x PBS. 700 μL of 2 mg/mL tissue lysates were added to 1.7 mL tubes for each biological replicate. 350 μL of each 2 mg/mL biological replicate for each sex was added to a protein quantitation master sample and 700 μL was aliquoted into 1.7 mL tubes for each treatment (probe, competition, and no probe). To competition samples, 2.8 μL each of 50 mM fentanyl hydrochloride and 50 mM remifentanil hydrochloride in 1:1: DMSO:H_2_O were added (200 μM each final concentration). To all probe positive and no probe control samples, an equivalent volume of DMSO:H_2_O vehicle control was added. The samples were vortexed briefly and incubated on the thermal mixing blocks at 500 rpm, 37 °C for 30 min. After the incubation with the competitor, 2.8 μL of 5 mM FA-T1 probe in DMSO was added to all probe positive and competition samples (20 μM FA-T1 final concentration). 3 μL of DMSO vehicle control were added to all no probe samples. The samples were vortexed and incubated at 500 rpm, 37 °C for 30 min. After probe incubation samples were transferred to 24 well plates and irradiated under UV light for 7 min. After irradiation the samples were transferred back to the corresponding 1.7 mL tubes and 8 μL from each biological replicate of the same sex and treatment were combined into a single 1.7 mL tube for SDS-PAGE gel analysis. Detailed methods for subsequent click chemistry attachment of biotin to probe-labeled proteins, enrichment on streptavidin agarose beads, on-bead tryptic digest, and tandem mass tag (TMT) labeling of peptides are described in the Supporting Information.

### LC-MS/MS acquisition

AfBPP samples were analyzed using a Waters nanoAcquity ultra performance liquid chromatography (UPLC) system (Milford, MA) connected to a Q-Exactive Plus Orbitrap mass spectrometer (Thermo Scientific, San Jose, CA). Samples were loaded into a precolumn (150 μm i.d., 4 cm length, packed in-lab with Jupiter C18 packing material, 300 Å pore size, 5 μm particle size; Phenomenex, Torrance, CA, USA) using mobile phase A (0.1% formic acid in water). The separation was carried out using a New Objective 75 µm i.d., 25-cm column with integrated emitter (Littleton, MA), packed in-lab with Waters BEH C18 1.7 µm packing material, (Milford, MA) at a flow rate of 200 nL/min, with column temperature of 45 °C. Mobile phases consisted of (A) 0.1% formic acid in water and (B) 0.1% formic acid in acetonitrile with the following gradient and column wash profile (min, %B): 0, 1; 10, 8; 105, 25; 115, 35; 120, 75; 123, 95; 129, 95; 130, 50; 132, 95; 138, 95; 140, 1. Data acquisition was started 20 min after sample load and inject to account for column dead volume and continued for 120 min with the remainder of the time used for column wash and column regeneration. The mass spectrometer source was set at 2.2 kV, and the ion transfer capillary was heated to 250 °C. The data-dependent acquisition mode was employed to automatically trigger the precursor scan and the MS/MS scans. The MS1 spectra were collected at a scan range of 300-2000 m/z, a resolution of 70,000, an automatic gain control (AGC) target of 3 × 10^6^, and a maximum injection ion injection time of 20 ms. For MS2, top 12 most intense precursors were isolated with a window of 0.7 m/z and fragmented by higher-energy collisional dissociation (HCD) with a normalized collision energy at 30%. The Orbitrap was used to collect MS/MS spectra at a resolution of 35,000, a maximum automated gain control (AGC) target of 1 × 10^5^, and maximum ion injection time of 100 ms. Each parent ion was fragmented once before being dynamically excluded for 30 s.

### Proteomics data analysis

MASIC (MS/MS Automated Selected Ion Chromatogram generator)^44^ and MSGF+^45^ (v2023.01.12) were used for peptide abundance and identification, as previously described. Reference protein collections were downloaded from UniProt on 2023-09-23 for *Homo sapiens, Macaca mulatta, Mustela furo, Cavia porcellus, Rattus norvegicus*, and *Mus musculus*. Peptide searches were performed for partially tryptic peptides with the following modifications: methionine oxidation (+15.9949), cysteine alkylation (+57.0215), and TMT-labeling of lysines and N-terminal peptides (+229.1629) with a parent ion tolerance of 20 ppm.

MASIC and MSGF+ results were aggregated using Mage software and a 5% false discovery rate (FDR) based on Q-value was used for initial filtering. Subsequent analyses were conducted using chemoprotR (https://code.emsl.pnl.gov/multiomics-analyses/chemoproteomicsR/-/tree/main). Peptide identifications for each organism were filtered using a MSGF spectral probability value of 3.29006 × 10^−9^ corresponding to a 1.9% FDR at the peptide level, calculated using a target-decoy approach.^46^ Once filtered, peptide redundancies were summed. Potential outliers were calculated using a robust Mahalanobis distance, calculated by using multiple metrics such as the correlation of samples to other samples in the same protocol, skewness, and MAD (median absolute distance) of the molecule abundance profile, and the proportion of missing values.^47^ Data were then normalized using mean centering^48, 49^ with a group-specific back transformation. Peptide-level data were rolled up to protein-level using a summation method. Reverse hits and contaminants were then removed from the dataset.

For quantitative analyses, an ANOVA was used to calculate fold changes between groups of interest and the corresponding p-values. For qualitative analyses, a G-test was conducted to identify significant differences between the proteins regarding presence/absence data. Proteins were determined to be statistically significant if they had a p-value < 0.05 and fold change (FC) > 1.5 for both probe only (ABP) vs. no probe (NP) and probe only (ABP) vs. fentanyl/remifentanil competition (Comp) comparisons.

### Homology mapping

Orthology relationships between the six model organisms of interest—human, rhesus macaque, ferret, guinea pig, rat, and mouse—were estimated by clustering proteins using three different algorithms. The first approach was Markov clustering (MCL), which is the least stringent, and returns clusters of proteins with similar function, but not necessarily precisely the same function. The second approach was reciprocal best match analysis, which clusters proteins based on best sequence similarity between organisms. This is a stringent approach, since it is constrained to attempt to identify only one member per organism, and it has a high probability of identifying functional orthologs across organisms. The final approach was OrthoFinder^50^, which performs a comprehensive phylogenetic analysis to identify true sequence orthologs. OrthoFinder clusters frequently have multiple members per organism due to gene duplication events. Results from the various clusterings were loaded into a relational database. This database was queried to identify orthologous proteins for all protein targets identified by FA-T1 chemoproteomic labeling, using the reciprocal best match results unless otherwise noted.

## Results

### Structure-based analysis of fentanyl AfBPP candidates

To evaluate fentanyl-AfBP designs prior to investing in their synthesis, we performed an *in silico* analysis to predict their properties and potential binding interactions using structure-based computational tools similar to our previous work.^51^ Molecular docking simulations were run for each fentanyl library compound with human MOR (PDB code 5C1M) to collect baseline scoring metrics. Fentanyl-AfBPs achieved a mean score of −8.9 (std. dev. ± 0.7) while decoys achieved a mean score of −8.3 (std. dev. ±0.9), a statistically significant difference. Fentanyl achieved a score of −9.2. Next, each candidate fentanyl probe was run through the docking simulation. Resulting docking scores suggested that the three proposed fentanyl-AfBPs will all mimic authentic fentanyl (**Figure 2**). Possible binding modes of the fentanyl-ABPs with MOR are depicted; each is juxtaposed with co-crystallized fentanyl experimentally determined using x-ray diffraction.

**Figure 2.**
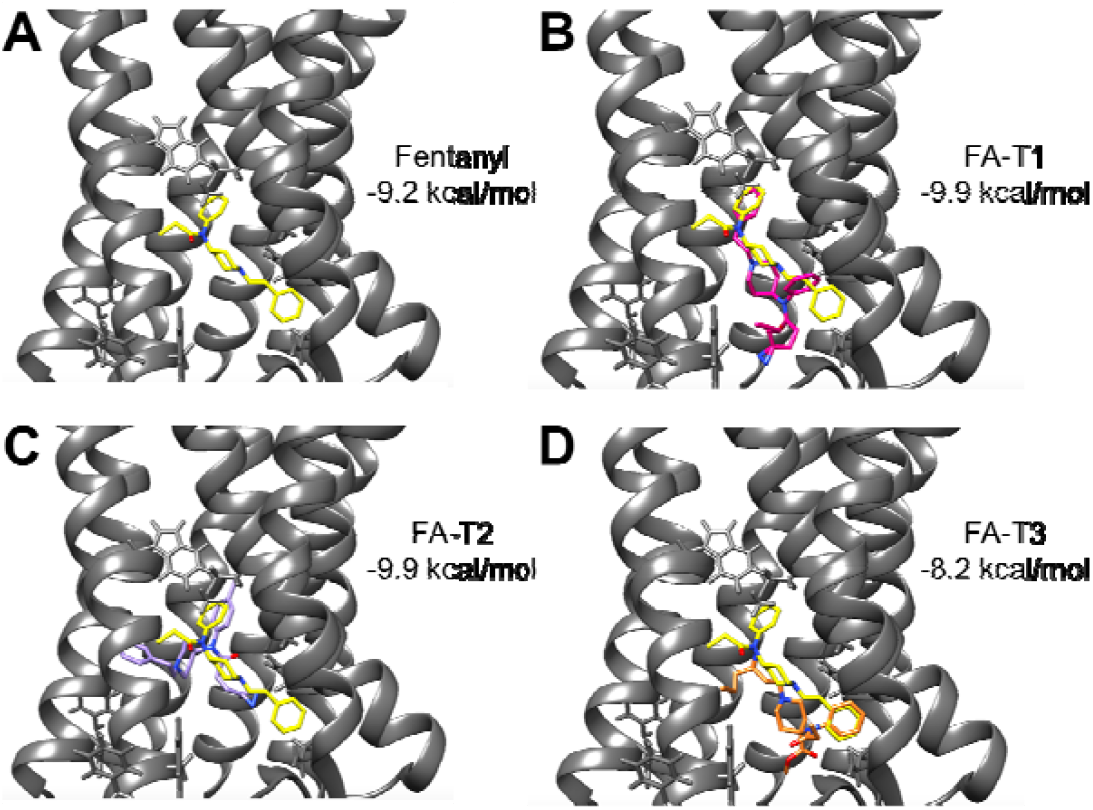
Predicted possible binding modes of the fentanyl-AfBPs at the fentanyl binding site of the mu opioid receptor (PDB 5C1M), shown in gray. Calculated MOR binding energies are displayed for each fentanyl analog. (A) Co-crystallized fentanyl, depicted in yellow. (B-D) AutoDock Vina predicted binding poses of FA-T1 (magenta), FA-T2 (lavender), and FA-T3 (orange) displayed alongside fentanyl.

### Synthesis and probe validation

Three fentanyl analog target (FA-T) probes were synthesized based on predicted favorable binding to MOR in docking simulations and general ease of the synthetic routes (**Figures S1-S3**). 4-anilino-N-phenethylpiperidine (4-ANPP) and 4-ANPP analogs were synthesized according to a previously published protocol.^52^ Initial attempts to perform amide coupling with 4-ANPP analogs using hexafluorophosphate azabenzotriazole tetramethyl uronium (HATU) failed. Amide coupling using Mukaiyama’s reagent was identified as an alternative for the synthesis of FA-T1 and FA-T2.^53, 54^ FA-T3 was synthesized in one step via alkylation of norcarfentanil.

### Probe labeling of tissues

Fluorescence imaging of SDS-PAGE gels showed that all fentanyl-AfBPs displayed dose-dependent labeling in rat tissue homogenates (**Figure S4**) The probes did not appear to label only the most abundant proteins, as determined from comparison with the total protein stain. While probe labeling could be discerned at concentrations as low as 5 µM in both rat brain and liver samples, labeling signal to noise compared to the no probe (NP) negative control was more robust at 10-20 µM of each probe. Notably, FA-T1 labeled proteins in rat brain homogenates more intensely than the other two probes, with FA-T3 having the weakest labeling (**Figure 3**). Treatment with a 10-fold excess of both fentanyl and remifentanil resulted in a slight reduction in FA-T1 labeling, suggesting competition for fentanyl-binding sites on proteins could be achieved under these conditions.

**Figure 3.**
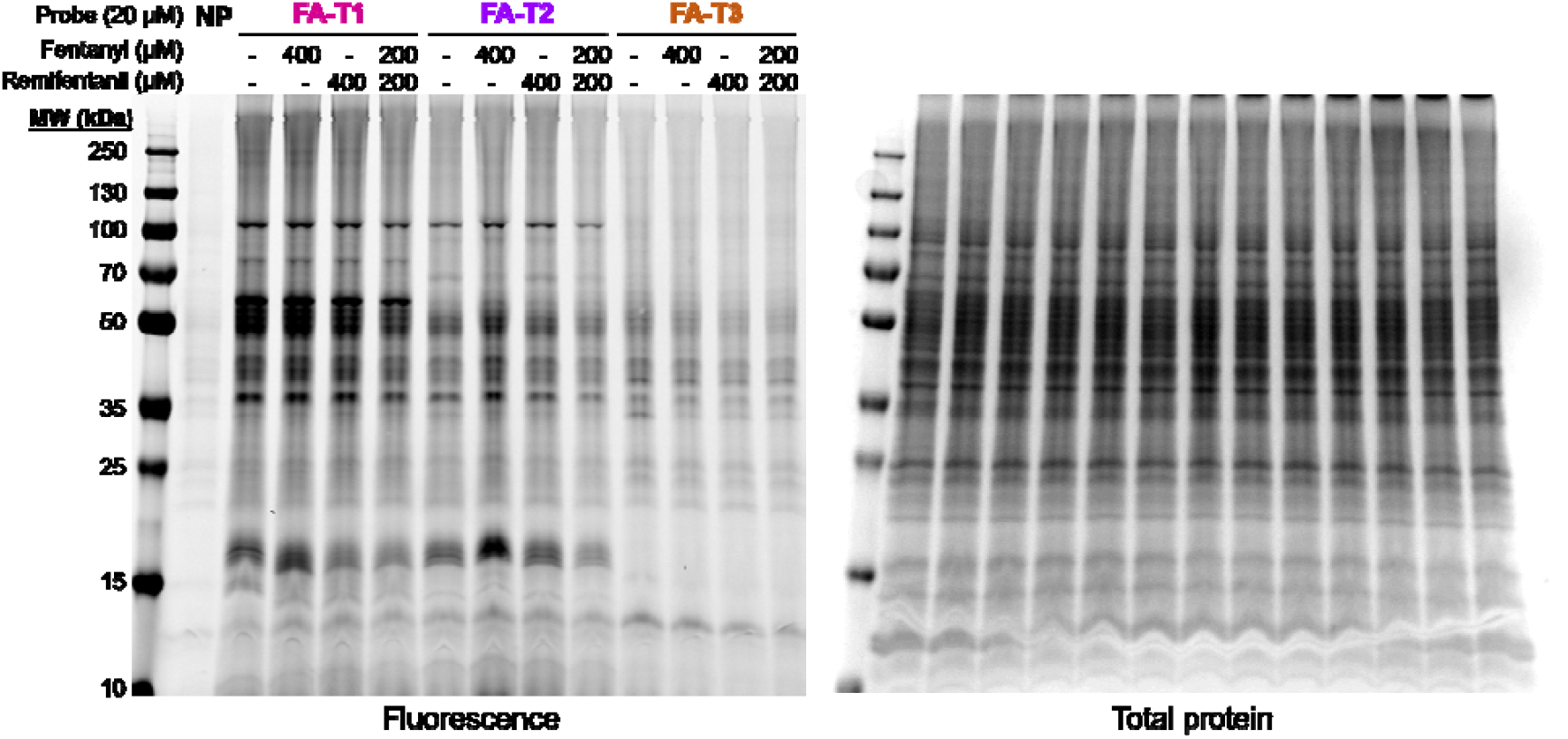
Labeling of rat brain homogenate by fentanyl probes (20 µM) with and without competition by fentanyl and/or remifentanil. No probe (NP) negative control sample labeled with vehicle (DMSO).

### Fentanyl protein target identification through AfBPP

Profiling of brain, heart, kidney, lung, and liver tissues from mouse, rat, guinea pig, ferret, rhesus macaque, and human yielded a list of proteins that were consistently identified across multiple organs and animal models. HNMT was the most frequently observed protein target of competitive AfBPP using fentanyl/remifentanil with the FA-T1 probe, being observed in 68% of all samples and in all profiled animal species (**Figure 4**). The next most frequently observed protein target, endophilin-B1 (SH3GLB1), was observed in more than half of all samples but was not observed as a statistically significant target of competitive AfBPP in human tissues. Fructosamine-3-kinase (FN3K) and CutA divalent cation tolerance homolog (CUTA) were observed in more than a third of all samples. FN3K is an ATP-dependent protein deglycation enzyme which may also have roles in mediating oxidative stress.^55, 56^ The role of CUTA in humans is not well understood, although studies in *E. coli* have shown it may play a role in small molecule signaling and transport.^57^ Studies of human CUTA show that it is distributed throughout the body and presumably plays a role in protein processing and trafficking.^58^ Competitive AfBPP with FA-T1 also identified the voltage-dependent anion channel proteins VDAC1 and VDAC2 in more than 20% of all datasets. VDAC proteins have previously been reported as preferred protein targets of diazirine-containing photoaffinity probes,^59^ although these proteins are also known to interact with other small molecule drugs that inhibit their oligomerization and prevent apoptosis.^60, 61^ Two monoamine oxidases, MAOB and MAOA, were also among the top 20 protein targets.

**Figure 4.**
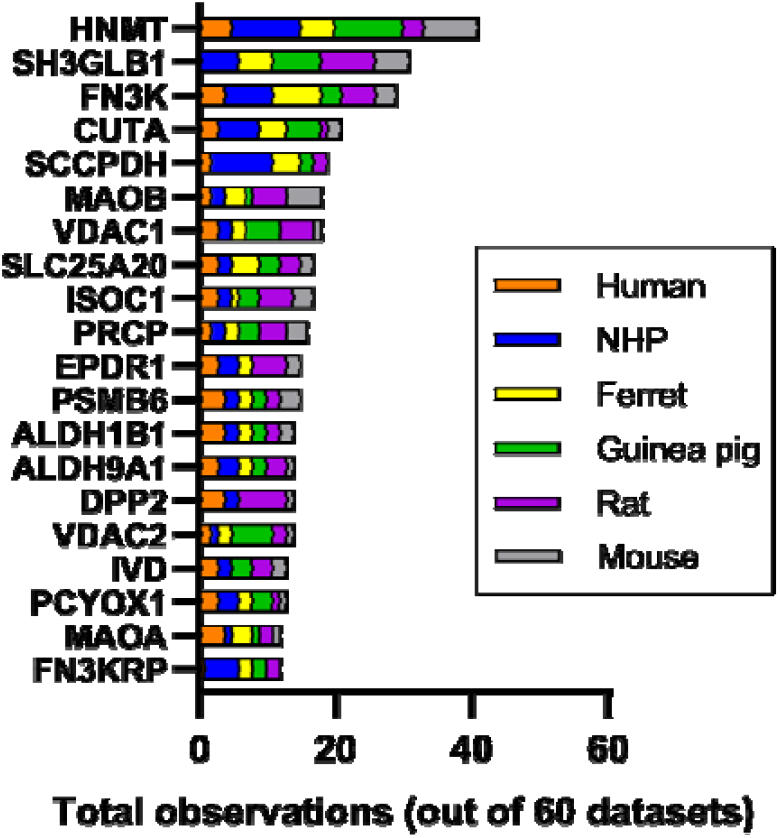
Top 20 observed protein targets of fentanyl (p < 0.05, FC > 1.5) identified using competitive AfBPP for fentanyl/remifentanil with the FA-T1 probe across all animal models, based on homology to a human protein. Data represents 5 tissues (brain, heart, kidney, lung, liver) in male and female organisms (n = 3 each), except for male human brain samples (n = 2), for a total of 60 datasets.

We did not identify MOR using AfBPP in any of the profiled samples, including brain tissues. MOR expression varies across cell types within tissues and species,^5^ and low expression levels may present a challenge for proteomic detection of this GPCR. We were also unable to detect MOR in rat brain tissue homogenates through Western blotting and global proteomics analysis. To enhance detection of membrane proteins by AfBPP, we processed rat brain tissues to obtain the membrane fraction. While we still did not identify MOR in these samples using FA-T1, we identified a larger number of membrane proteins (**Table 1**), including the muscarinic acetylcholine receptor subtype M1 (CHRM1) and three transporters (MFSD4B2, SLC6A1, SLC25A20) among the top proteins with largest average fold change for fentanyl/remifentanil competition in both male and female groups. Several proteins previously identified in whole tissue homogenate experiments were also detected in these samples, including CUTA, FN3K, DPP2, SLC25A20, VDAC1, PCYOX1, SH3GLB1, and SCCPDH.

**Table 1.** Selected protein targets identified in rat membrane fraction proteomes from competitive AfBPP using fentanyl/remifentanil and FA-T1. Proteins shown are the top 25 proteins with p-value < 0.05 having the highest mean fold change (FC) for female and male sample groups (n = 3).

| UniProtID | Protein | Protein description | Male FC | Female FC |
| --- | --- | --- | --- | --- |
| Q6MGD0 | CUTA | Protein CutA | 4.68 | 7.50 |
| Q6IG04 | K2C72 | Keratin, type II cytoskeletal 72 | 6.25 | 4.75 |
| P03889 | MTND1 | NADH-ubiquinone oxidoreductase chain 1 | 6.42 | 2.60 |
| Q6IG02 | K22E | Keratin, type II cytoskeletal 2 epidermal | 6.82 | 2.14 |
| D3ZZU8 | FN3K | Fructosamine 3-kinase | 3.03 | 4.11 |
| D3Z9X5 | MFSD4B2 | Similar to Na<sup>+</sup> dependent glucose transporter 1 | 2.32 | 3.93 |
| P23978 | SLC6A1 | Sodium- and chloride-dependent GABA transporter 1 | 3.10 | 2.98 |
| Q9EPB1 | DPP2 | Dipeptidyl peptidase 2 | 2.63 | 3.23 |
| P97521 | SLC25A20 | Mitochondrial carnitine/acylcarnitine carrier protein | 2.17 | 3.46 |
| P08482 | CHRM1 | Muscarinic acetylcholine receptor M1 | 2.00 | 3.63 |
| Q63524 | TMED2 | Transmembrane emp24 domain-containing protein 2 | 2.69 | 2.73 |
| P49655 | KCNJ10 | ATP-sensitive inward rectifier potassium channel 10 | 2.70 | 2.69 |
| P00406 | MT-CO2 | Cytochrome c oxidase subunit 2 | 2.79 | 2.52 |
| A0A0G2JXD0 | RPL9L2 | Large ribosomal subunit protein uL6 | 3.21 | 2.07 |
| P52555 | ERP29 | Endoplasmic reticulum resident protein 29 | 3.00 | 2.25 |
| Q9Z2L0 | VDAC1 | Voltage-dependent anion-selective channel protein 1 | 2.36 | 2.82 |
| Q99ML5 | PCYOX1 | Prenylcysteine oxidase 1 | 2.35 | 2.82 |
| B5DF51 | MMGT1 | ER membrane protein complex subunit 5 | 2.65 | 2.46 |
| Q62636 | RAP1B | Ras-related protein Rap-1b | 3.08 | 1.91 |
| P35332 | HPCAL4 | Hippocalcin-like protein 4 | 2.31 | 2.55 |
| P20070 | CYB5R3 | NADH-cytochrome b5 reductase 3 | 2.32 | 2.53 |
| P19944 | RPLP1 | Large ribosomal subunit protein P1 | 3.19 | 1.57 |
| P25113 | PGAM1 | Phosphoglycerate mutase 1 | 2.46 | 2.28 |
| Q6AYE2 | SH3GLB1 | Endophilin-B1 | 1.77 | 2.93 |
| Q6AY30 | SCCPDH | Saccharopine dehydrogenase-like oxidoreductase | 1.82 | 2.87 |

### Docking simulations to assess fentanyl and remifentanil binding with selected protein targets

To further evaluate the experimental AfBPP binding results, we conducted docking simulations for 6 scenarios (i.e., fentanyl and remifentanil binding to HNMT, MAOB, and CHRM1, combinatorially) (**Table 2, Figure 5-6, Figure S5**). These results show likely binding modes corroborating our AfBPP results *in silico*.

**Table 2.** Binding energies of fentanyl and remifentanil to protein targets identified by competitive AfBPP.

| Protein | Ligand | Binding energy* |
| --- | --- | --- |
| HNMT<br>(PDB ID: 2AOT) | Fentanyl | -10.2 |
|  | Remifentanyl | -7.0 |
| MAOB<br>(PDB ID: 6FW0) | Fentanyl | -10.4 |
|  | Remifentanyl | -9.4 |
| CHRM1<br>(PDB ID: 6ZFZ) | Fentanyl | -10.0 |
|  | Remifentanyl | -7.1 |

\* Unit is kcal/mol. A more negative value indicates stronger binding between the receptor and ligand.

**Figure 5.**
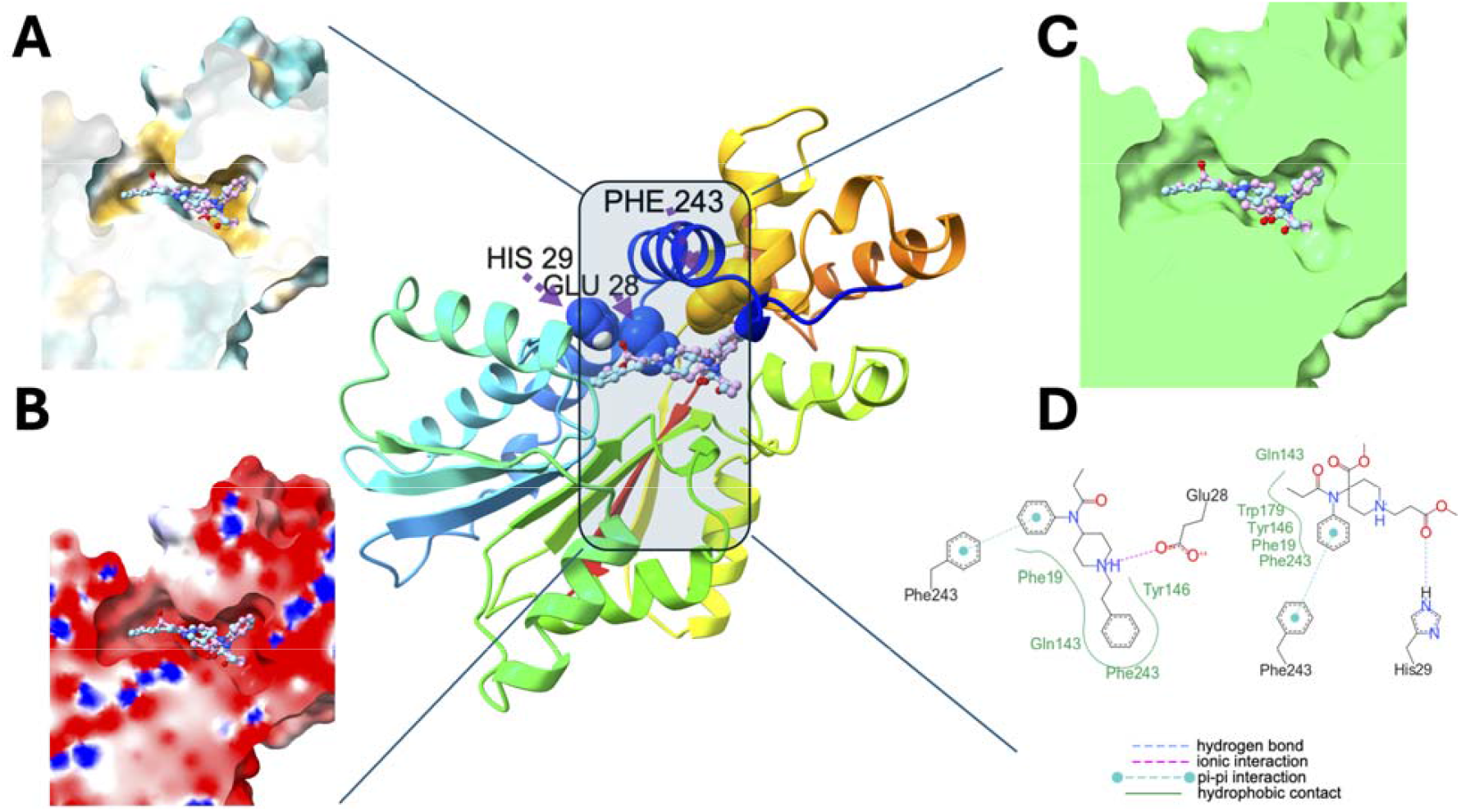
Docking simulation of fentanyl (cyan colored ball and stick) and remifentanil (pink colored ball and stick) binding into HNMT. Key interaction residues are rendered as spheres. (A) Putative binding pocket with hydrophobicity rendered. Cyan: hydrophilic region, white: intermediate lipophilicity region, goldenrod: lipophilic region. (B) Same view as A, but with electrostatic potential visualization. Blue: positive, white: neutral, red: negative. (C) Ligands show probable shape complementarity into binding pocket. (D) Fentanyl binding orientation is driven by favorable ionic interaction (i.e., between cation N6 of fentanyl and anion oxygen of Glu28 of HNMT) and pi-pi interaction (i.e., between fentanyl benzene ring and Phe243). The binding is further strengthened by hydrophobic contacts between Phe19, Gln143, Phe243 and Tyr146 of HNMT and fentanyl. Remifentanil binding orientation is driven by favorable hydrogen bond between remifentanil and His29 of HNMT and pi-pi interaction. The binding is further strengthened by hydrophobic contacts between remifentanil and HNMT.

**Figure 6.**
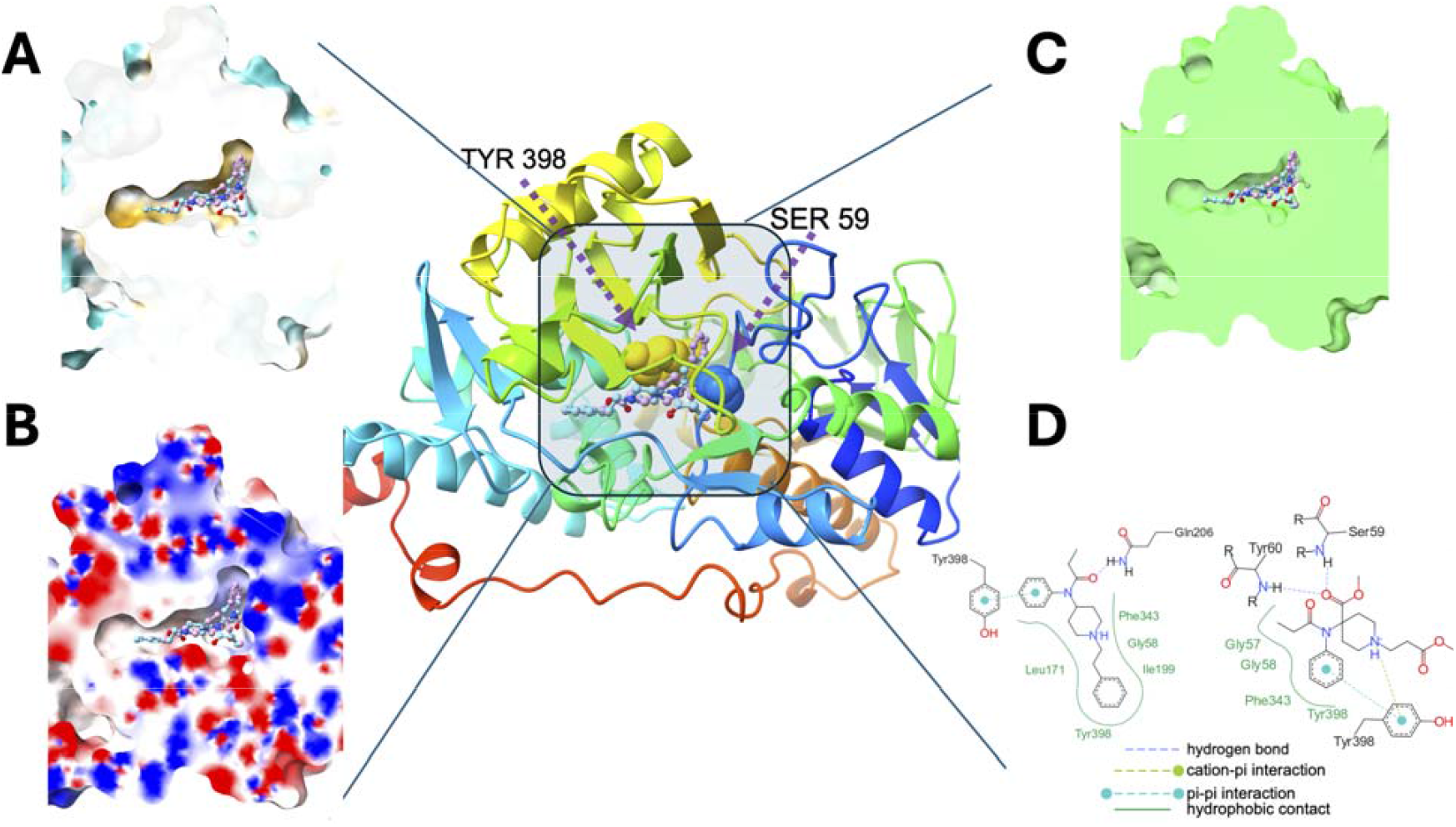
Docking simulation of fentanyl (cyan colored ball and stick) and remifentanil (pink colored ball and stick) binding into MAOB. Key interaction residues are rendered as spheres. (A) Putative binding pocket with hydrophobicity rendered. Cyan: hydrophilic region, white: intermediate lipophilicity region, goldenrod: lipophilic region. (B) Same view as A, but with electrostatic potential visualization. Blue: positive, white: neutral, red: negative. (C) Ligands show probable shape complementarity into binding pocket. (D) Fentanyl binding orientation is driven by favorable pi-pi interaction and hydrophobic interaction in large area. Remifentanil binding orientation is driven by pi-pi interaction, cation-pi interaction, hydrogen bonds and hydrophobic contacts.

We found that computational modeling of AfBPP results improved our ability to prioritize protein targets for further experimental analyses with its calculated binding energies and manual inspection. For example, most of the top docking poses consistently converged on the same binding pockets known for experimentally observed ligand positions in each modeled protein target. This suggests that the ligand binding identified by our AfBPP experiment is more likely stable rather than nonspecific or spurious. Although we used experimentally determined receptor structures for docking, the corresponding AlphaFold-predicted models demonstrate that these protein receptors are quite stable.^62^ This suggests that the selections from our AfBPP experiment analysis are based on well-folded proteins rather than intrinsically disordered ones. As a note, while AutoDock Vina^63^ is known to provide slightly less accurate binding energies compared to AutoDock4, it generally produces more reliable binding poses.^64^ Given the alignment of the AutoDock series with experimental results,^63^ we believe our trend analysis remains robust and provides valuable insights.

## Discussion

Our fundamental understanding of fentanyl’s biological effects has been dominated by its agonism of opioid receptors, particularly MOR. However, we have limited data on fentanyl binding interactions with proteins beyond opioid receptors. Thus, our primary goal for design and validation of fentanyl-based AfBPs for non-opioid receptor target identification was to confirm the ability of proteins to recognize the AfBPs in a similar manner as fentanyl. Of the three AfBPs, FA-T1 displayed the highest protein labeling intensities in rat brain homogenates in gel analyses. Previous work by the Janda group on fentanyl hapten design for immunotherapy indicated that modification of the amide produces antibodies that recognize fentanyl.^65^ Based on our experimental results and literature precedent, we selected FA-T1 for further experiments.

Fentanyl’s high lipid solubility has been proposed as a key factor affecting its pharmacological profile, including its enhanced ability to cross the blood-brain barrier and slower clearance compared to more hydrophilic opioids, such as morphine.^66^ However, the lipophilicity of fentanyl and its analogs poses a significant challenge to its application for chemoproteomic profiling due to non-specific, hydrophobic interaction. We required a relatively high concentration of fentanyl-AfBPs (20 µM) to achieve sufficient labeling of proteins in our tissue homogenates for robust identification. We surmise that the composition of these samples, which were centrifuged at relatively low speeds and included pieces of the cellular membrane in suspension, may favor probe partitioning into lipid environments; attempting AfBPP with lower probe concentrations (1 µM) resulted in poor protein identifications.

Our broad profiling study across multiple tissue types and animal species sought to identify non-opioid receptor protein targets of fentanyl analogs that may contribute to fentanyl’s toxicity throughout the body. By performing pre-incubation of samples with authentic fentanyl and remifentanil, we attempted to identify specific proteins that bind to actual fentanyl analogs. Notably, the numbers of significant proteins identified through this approach was substantially lower than probe compared to no probe (NP) samples alone, suggesting this competitive AfBPP method is critical for differentiating more selective protein targets over non-specific binding interactions. We have also previously used this strategy to identify proteins that bind the hydrophobic lipid benzoquinone, sorgoleone, in bacterial lysates.^67^

Consistent with previous reports that fentanyl can bind to muscarinic receptors in rat brain at high doses,^15^ we identified the M1 subtype through AfBPP in rat brain membrane fraction. Previous studies observed antimuscarinic activity during high dose fentanyl treatment,^16^ with the M3 acetylcholine muscarinic receptor subtype confirmed to bind fentanyl in the heart for rat and porcine models,^17^ although Hustveit noted no apparent muscarinic receptor subtype selectivity in rat brain. While profiling unfractionated tissue homogenates identified multiple membrane proteins such as CUTA, SH3GLB1, VDAC1, and SLC25A20, specific analysis of the membrane fraction yielded more consistent identification of membrane proteins that were not as frequently observed in whole tissue homogenates, such as SLC6A1, which was recently reported to have increased expression in midbrain organoid cells exposed to fentanyl.^68^ Expanded AfBPP of fractionated or region-specific proteomes in the future may improve sensitivity for proteins that are concentrated in specific cellular regions or cell types.

We most frequently identified the cytosolic histamine-degrading enzyme HNMT using FA-T1 labeling with fentanyl and remifentanil competition across whole tissue homogenates from all animal species. HNMT is expressed throughout the body, including in the kidney, liver, lung, heart, and central nervous system.^69, 70^ HNMT was shown to bind tightly to other drugs containing aromatic groups such as diphenhydramine, amodiaquine, metoprine, and tacrine, resulting in enzyme inhibition,^71^ although HNMT’s potential interaction with fentanyl analogs has not been previously reported to date. Histamine release is known to be a significant side effect of opioids such as morphine,^72^, but fentanyl generally has not been reported to cause increases in histamine levels.^73^ However, a 2013 study observed increased histamine in bronchoalveolar lavage fluid (BALF) from fentanyl-exposed mice,^74^ and antihistamines have been shown to reduce fentanyl-induced cough.^75^ Interestingly, we also identified both MAOA and MAOB as protein targets for all animal species in our whole tissue AfBPP study. Monoamine oxidases (MAOs) are involved in the catabolism of monoamines, including neurotransmitters such as histamine.^76^ MAOB preferentially oxidizes 1-methylhistamine, the product of HNMT action upon histamine, but may also oxidase histamine itself when HNMT is inhibited.^77^ Whether inhibition of HNMT and MAOs by fentanyl could contribute to increased histamine or otherwise perturb signaling of this neurotransmitter in different tissues remains to be determined.

In summary, we generated a fentanyl-based AfBP, FA-T1, that was applied to identifying protein targets of fentanyl analogs across multiple tissue types and animal models using a competitive AfBPP approach. Combined with computational modeling of protein-fentanyl modeling, this chemoproteomic strategy enables broad screening for proteins of interest in complex proteomes to elucidate potential protein interactions that may be responsible for fentanyl’s adverse effects. In this work, a single high concentration of equimolar fentanyl and remifentanil was used to compete for protein binding sites with FA-T1, and therefore all proteins identified likely represent targets with greater relevance to high dose exposures such as those seen in overdose scenarios rather than lower therapeutic doses. Future studies investigating a range of competitor concentrations may enable differentiation of protein targets that bind fentanyl analogs at low vs. high concentrations. Our comprehensive profiling of five tissue types across six mammalian species, including humans, provides new insights into protein binding interactions that may help to elucidate mechanisms of toxicity for fentanyl.

## Acknowledgements

This work was supported by the Defense Threat Reduction Agency Joint Science and Technology Office (DTRA-JSTO) for Chemical and Biological Defense under Grant Numbers HDTRA1033013, HDTRA1343479, and HDTRA1343354. We would like to thank Heather Olson, Carrie Nicora, and Marina Gritsenko for sample preparation advice. We are grateful to Wendy Price and staff at the Oregon National Primate Research Center (ONPRC) Tissue Distribution Program for providing the rhesus macaque tissues for this work. The collection of non-human primate tissues used in this research was supported in part by National Institutes of Health Grant P51OD011092 to the Oregon National Primate Research Center. A portion of this research was supported by the EMSL user project award 51814 (https://dx.doi.org/10.46936/reso.proj.2021.51814/60000335), for leveraging instrument capabilities operating at the Environmental Molecular Sciences Laboratory, a DOE Office of Science User Facility. PNNL is operated by Battelle for the DOE under contract DE-AC05-76RL01830. Figure 1 was generated in part using BioRender.

