## Supporting Information for "Discovery of non-opioid receptor protein targets of fentanyl and remifentanil by affinity-based protein profiling in diverse animal model and human tissues"

*Corresponding authors:

**General synthetic methods**

To reduce exposure risks to research personnel during work with fentanyl analogs,^1^ synthesis and handling of fentanyl analogs in pure or solid form was performed in a ventilated enclosure with HEPA filtration with appropriate personal protective equipment (PPE), including chemically resistant long sleeve aprons, double nitrile gloves with inner gloves secured to apron sleeves with chemically resistant seam-sealing tape, and cleanroom sleeves.

Chemicals were purchased from commercial suppliers such as Acros, Alfa Aesar, TCI America, Fisher, VWR, and Sigma-Aldrich and used without further purification unless otherwise noted. 3-(But-3-yn-1-yl)-3-(2-iodoethyl)-3H-diazirine was purchased from Ambeed. 3-(3-(But-3-yn-1-yl)-3H-diazirin-3-yl)propanoic acid was purchased from Sigma-Aldrich.

Dry acetonitrile, dimethylformamide (DMF), tetrahydrofuran (THF), and dichloromethane (CH_2_Cl_2_) were obtained from an LC Tech SP-1 Stand Alone Solvent Purification System. Automated flash silica gel column chromatography was performed using a Biotage Isolera purification system. Analytical thin layer chromatography (TLC) was performed using silica gel 60 F254 plates (0.25 mm) and compounds visualized by shortwave UV irradiation with a handheld UV lamp or staining as indicated.

^1^H and ^13^C NMR spectra of synthetic intermediates were acquired in CDCl_3_ (Cambridge Isotopes, Tewksbury, MA) at ambient temperature (25 °C) on a Bruker 400 MHz Avance III spectrometer equipped with a 5 mm BBFO SmartProbe. Fentanyl analogs were analyzed using an Agilent Inova 500 MHz NMR. All chemical shifts are reported in the standard notation of parts per million the peak of the residual proton or carbon signal of CDCl_3_ (^1^H NMR δ 7.26 and ^13^C NMR δ 77.36 ppm) as an internal reference. Splitting patterns are indicated as follows: s, singlet; d, doublet; t, triplet; m, multiplet; dd, doublet of doublets. Low resolution mass spectrometry (LRMS) with electrospray ionization (ESI) was performed using a Finnigan LTQ mass spectrometer (Thermo Electron Corporation).

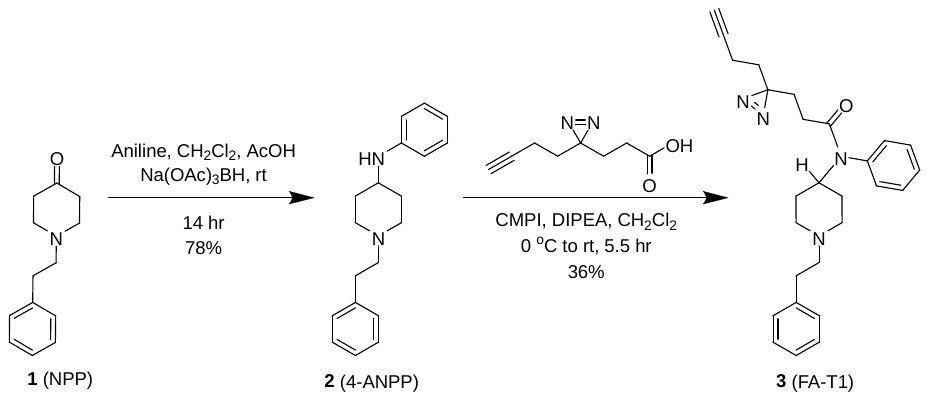

**Figure S1.** Fentanyl probe FA-T1 synthetic scheme.

4-anilino-N-phenethylpiperidine (compound 2; 4-ANPP)^2^

Aniline (0.209 g, 2.2 mmol, 1 equiv) was dissolved in 10 mL CH_2_Cl_2_ in a 50 mL round bottom flask with stirbar. The resulting solution was cooled in an ice bath, and acetic acid (0.12 mL, 2.2 mmol, 1 equiv) was added dropwise. To the reaction mixture, N-phenylethylpiperidin-4-one (NPP, **1**; 0.458 g, 2.2 mmol, 1 equiv) dissolved in 3 mL CH_2_Cl_2_ was added gradually. Then, sodium triacetoxyborohydride (0.714 g, 3.4 mmol, 1.5 equiv) was added, portionwise. The reaction was stirred, warming to r.t. overnight. After ~16 hr, 5 mL MeOH was added to quench the reaction. The crude reaction mixture was transferred to a separatory funnel, diluted with 100 mL CH_2_Cl_2_, washed with sat. aq. NaHCO_3_, and dried over Na_2_SO_4_. The crude product was concentrated in vacuo to a light yellow oil that solidified to a pale yellow solid upon standing. The product was purified by flash column chromatography, eluting 50-90% EtOAc/hex. The combined product fractions (R_f_ = 0.17, 50% EtOAc/hex) were dried by rotary evaporation to a white crystalline solid (0.494 g, 78.5% yield). ^1^H NMR (400 MHz, CDCl_3_) δ 7.31 – 7.22 (m, 2H), 7.18 (dd, *J* = 14.2, 7.3 Hz, 4H), 7.17 – 7.10 (m, 1H), 6.71 – 6.62 (m, 1H), 6.62 – 6.55 (m, 2H), 3.55 (s, 1H), 3.31 (tt, *J* = 9.9, 3.9 Hz, 1H), 3.01 – 2.91 (m, 2H), 2.86 – 2.77 (m, 2H), 2.66 – 2.57 (m, 2H), 2.26 – 2.16 (m, 2H), 2.12 – 2.03 (m, 2H), 1.58 – 1.44 (m, 2H).

**3-(3-(but-3-yn-1-yl)-3H-diazirin-3-yl)-N-(1-phenethylpiperidin-4-yl)-N-phenylpropanamide (3, FA-T1)**

3-(3-(But-3-yn-1-yl)-3H-diazirin-3-yl)propanoic acid (0.0455 g, 0.27 mmol, 1 equiv) and 2-chloro-1-methylpyridinium iodide (CMPI; 0.115 g, 0.45 mmol, 1.6 equiv) were added to an oven-dried 50 mL round bottom flask with stirbar. The flask was flushed with N_2_, and 1 mL dry CH_2_Cl_2_ was added. Intermediate **2** (4-ANPP; 0.0986 g, 0.35 mmol, 1.3 equiv) was dissolved in 2 mL dry CH_2_Cl_2_ and then added to the reaction flask. A balloon of argon gas was attached to the flask, and the reaction was cooled over an ice bath. DIPEA (0.26 mL, 1.5 mmol, 5.5 equiv) was added to the reaction, dropwise. The reaction turned a cloudy, light yellow color. The ice bath was removed, and the reaction was stirred at r.t. for 5.5 hr. The reaction was quenched by adding water (20 mL), and the product was extracted with 3x CH_2_Cl_2_. The combined organic layers were washed with NaCl brine (20 mL) and dried over Na_2_SO_4_. The crude product was dried via rotary evaporation to afford a red residue. The product was purified by column chromatography (12 g SiO_2_), eluting 25 to 50 to 80% EtOAc/hex (Rf = 0.24, 2:1 EtOAc/hex). The fractions containing the product **3** were combined and dried to a clear residue by rotary evaporation (0.0424 g, 36.1% yield). NMR characterization is presented in Table S1. LRMS-ESI (m/z): calcd for C_27_H_32_N_4_OH^+^ [M+H^+^]: 429.27. Found 429.30.

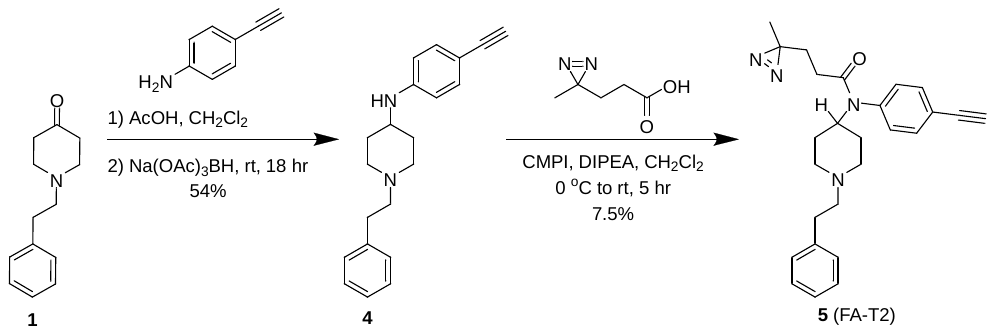

**Figure S2.** Fentanyl probe FA-T2 synthetic scheme.

**N-(4-ethynylphenyl)-1-phenethylpiperidin-4-amine (4)
4** was synthesized using reductive amination conditions. 4-ethynylaniline (0.269 g, 2.3 mmol, 1 equiv) and NPP (**1**; 0.468 g, 2.3 mmol, 1 equiv) were added to an oven-dried, 2-neck round bottom flask with stirbar. The flask was placed under a nitrogen atmosphere, and CH_2_Cl_2_ (10 mL) and acetic acid (0.13 mL, 2.3 mmol, 1 equiv) were added. The flask was cooled over an ice bath. Solid Na(OAc)_3_BH (0.726 g, 3.4 mmol, 1.5 equiv) was added, portion-wise, to the flask while stirring. After full addiction of the reducing agent, the flask was stirred overnight under nitrogen, warming to r.t. After ~18 hr, the reaction was quenched with 5 mL MeOH, diluted with 50 mL CH_2_Cl_2_, and transferred to a separatory funnel. The product was washed with sat. aq. NaHCO_3_ and then extracted with 2x CH_2_Cl_2_. The combined organic layers were washed with NaCl brine and dried over Na_2_SO_4_. The crude product was concentrated in vacuo to an amber oil and purified by silica gel flash chromatography, eluting 50-80% EtOAc/hex (product R_f_ = 0.19, 80% EtOAc/hex). The product **4** was isolated as a white crystalline solid (0.376 g, 54% yield). ^1^H NMR (400 MHz, CDCl_3_) δ 7.38 – 7.29 (m, 4H), 7.25 (d, *J* = 7.4 Hz, 3H), 6.56 – 6.49 (m, 2H), 3.80 (d, *J* = 8.0 Hz, 1H), 3.33 (dtt, *J* = 9.8, 6.3, 3.7 Hz, 1H), 2.98 (d, *J* = 17.8 Hz, 3H), 2.90 – 2.81 (m, 2H), 2.69 – 2.61 (m, 2H), 2.22 (td, *J* = 11.4, 2.6 Hz, 2H), 2.13 – 2.04 (m, 2H), 1.53 (dtd, *J* = 13.9, 10.6, 3.7 Hz, 2H). ^13^C NMR (101 MHz, CDCl_3_) δ 147.39, 140.29, 133.46, 128.67, 128.38, 126.04, 112.54, 109.56, 84.78, 74.83, 60.49, 52.26, 49.57, 33.82, 32.29. LRMS-ESI (m/z): calcd for C_21_H_24_N_2_H^+^ [M+H^+^]: 305.20. Found 305.26.

**N-(4-ethynylphenyl)-3-(3-methyl-3H-diazirin-3-yl)-N-(1-phenethylpiperidin-4-yl)propenamide (5, FA-T2)**

To a 50 mL oven-dried round bottom flask with stirbar, CMPI (0.185 g, 0.72 mmol, 1.45 equiv) and 3-(3-methyl-3H-diazirin-3-yl)propanoic acid (0.064 g, 0.50 mmol, 1 equiv) were dissolved in 5 mL dry CH_2_Cl_2_. The reaction mixture was stirred, forming a yellow solution with suspended solids. An argon balloon was attached to the reaction flask. The reaction mixture was cooled in an ice water bath and dry DIPEA (0.4 mL, 2.5 mmol, 5 equiv) was added, dropwise. A solution of alkyne intermediate **4** (0.140 g, 0.46 mmol, 0.92 equiv) dissolved in 1 mL CH_2_Cl_2_ was added to the reaction mixture, dropwise. The flask was removed from the ice water bath and stirred at r.t. for 5 hr. The reaction was quenched by adding 20 mL water. The product was extracted with 3x CH_2_Cl_2_, and the combined organic layers were washed with NaCl brine (20 mL) and dried over Na_2_SO_4_. The crude product was dried by rotary evaporation to an orange-brown residue. The product was purified by flash column chromatography, equilibrating a 12 g SiO_2_ prepacked column with 40% EtOAc/hex and eluting with 40-50% EtOAc/hex. The product was isolated as a clear residue (14.3 mg, 7.5% yield). NMR characterization is presented in Table S1. LRMS-ESI (m/z): C_26_H_30_N_4_OH^+^ calcd for H^+^ [M+H^+^]: 415.25. Found 415.22

**Methyl 1-(2-(3-(but-3-yn-1-yl)-3H-diazirin-3-yl)ethyl)-4-(N-phenylpropionamido)piperidine-4-carboxylate (7, FA-T3)**
Norcarfentanil (**6**; 0.975 g, 0.34 mmol, 1 equiv) was weighed into an oven-dried round bottom flask and placed under an argon atmosphere. Anhydrous acetonitrile (5 mL) was added to dissolve the starting material. The reaction mixture was then cooled in an ice bath. (3-(But-3-yn-1-yl)-3-(2-iodoethyl)-3H-diazirine (0.06 mL, 0.39 mmol, 1.15 equiv) dissolved in 1 mL dry acetonitrile was added to the reaction mixture. The ice bath was then removed, and the reaction was stirred at r.t. for 5.5 hr. The reaction was quenched with 25 mL water and diluted with CH_2_Cl_2_. The product was extracted with 3x CH_2_Cl_2_. The combined organic layers were washed with NaCl brine and dried over Na_2_SO_4_. The crude reaction mixture was concentrated by rotary evaporation, and the crude product was purified manually by flash chromatography on a 12 g SiO_2_ column, eluting 0-5% MeOH/ CH_2_Cl_2_ (R_f_ = 0.50, 50% MeOH/CH_2_Cl_2_). The product **7** was isolated as a light amber residue (88 mg, 64% yield). NMR characterization is presented in Table S1. LRMS-ESI (m/z): calcd for C_23_H_30_N_4_O_3_H^+^ [M+H^+^]: 411.23. Found 411.24.

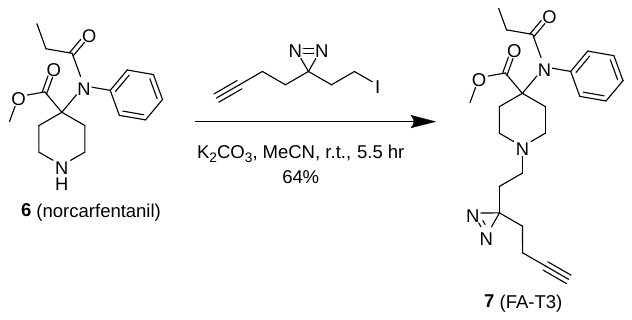

**Figure S3.** Fentanyl probe FA-T3 synthetic scheme.

**Table S1.**	Chemical shifts for fentanyl and fentanyl probes FA-T1, FA-T2, and FA-T3.

|  | **Fentanyl** | | **FA-T1** | | **FA-T2** | | **FA-T3** | |
| --- | --- | --- | --- | --- | --- | --- | --- | --- |
| **Moiety** | ^1^H | ^13^C | ^1^H | ^13^C | ^1^H | ^13^C | ^1^H | ^13^C |
| ***Piperidine*** | | | | | | | | |
| 2 | 4.69 | 52.1 | 4.65 | 52.4 | 4.65 | 52.5 | -- | 62.7 |
| 3 | 1.44  1.81 | 30.6 | 1.43  1.80 | 30.5 | 1.41  1.79 | 30.4 | 1.61  2.26 | 33.4 |
| 4 | 2.16  3.00 | 53.1 | 2.15  3.00 | 50.3 | 2.15  3.00 | 53.0 | 2.36  2.58 | 49.8 |
| ***Methyl carboxylate*** | | | | | | | | |
| C=O | -- | -- | -- | -- | -- | -- | -- | 174.0 |
| OCH3 | -- | -- | -- | -- | -- | -- | 3.79 | 52.2 |
| ***Piperidine N-2-phenylethyl*** | | | | | | | | |
| 1 | 2.53 | 60.5 | 2.53 | 60.5 | 2.54 | 60.4 | 2.17 | 52.4 |
| 2 | 2.73 | 33.8 | 2.72 | 33.9 | 2.73 | 33.8 | 1.51 | 30.5 |
| 3 (aziridine) | -- | -- | -- | -- | -- | -- | -- | 27.2 |
| 4 | -- | -- | -- | -- | -- | -- | 1.60 | 32.5 |
| 5 | -- | -- | -- | -- | -- | -- | 1.98 | 13.3 |
| 6 (alkyne) | -- | -- | -- | -- | -- | -- | -- | 82.8 |
| 7 (alkyne) | -- | -- | -- | -- | -- | -- | 1.97 | 69.1 |
| *i* | -- | 140.2 | -- | -- | -- | 140.0 | -- | -- |
| *o* | 7.15 | 128.6 | -- | -- | 7.17 | 128.5 | -- | -- |
| *m* | 7.26 | 128.4 | -- | -- | 7.28 | 128.3 | -- | -- |
| *p* | 7.18 | 126.0 | -- | -- | 7.20 | 126.0 | -- | -- |
| ***Propanamide*** | | | | | | | | |
| 1 C=O | -- | 173.6 | -- | 171.0 | -- | 170.6 | -- | 174.1 |
| 2 | 1.93 | 28.5 | 1.67 | 29.1 | 1.26 | 29.7 | 1.88 | 29.1 |
| 3 | 1.02 | 9.6 | 1.75 | 28.0 | 1.70 | 29.4 | 0.96 | 9.2 |
| 4 (aziridine) | -- | -- | -- | 27.9 | -- | 25.5 | -- | -- |
| 5 | -- | -- | 1.56 | 32.5 | 0.94 | 20.0 | -- | -- |
| 6 | -- | -- | 1.94 | 13.2 | -- | -- | -- | -- |
| 7 (alkyne) | -- | -- | -- | 82.8 | -- | -- | -- | -- |
| 8 (alkyne) | -- | -- | 1.92 | 69.0 | -- | -- | -- | -- |
| ***Amide N-phenyl*** | | | | | | | | |
| *i* | -- | 138.8 | -- | 138.1 | -- | 138.7 | -- | 139.3 |
| *o* | 7.08 | 130.4 | 7.08 | 130.2 | 7.06 | 130.3 | 7.31 | 130.6 |
| *m* | 7.39 | 129.3 | 7.41 | 129.4 | 7.55 | 133.1 | 7.42 | 129.2 |
| *p* | 7.36 | 128.3 | 7.40 | 128.5 | -- | 122.6 | 7.41 | 128.7 |
| 1’ (alkyne) | -- | -- | -- | -- | -- | 82.4 | -- | -- |
| 2’ (alkyne) | -- | -- | -- | -- | 3.17 | 78.8 | -- | -- |

Tissue procurement

Brain, heart, lung, kidney, and liver tissues for mouse (Albino, Swiss Webster), rat (Sprague Dawley), guinea pig (Dunkin Hartley), ferret, and human were acquired from BioChemed (Winchester, Virginia). Non-human primate (Rhesus macaque) tissues were acquired from the Oregon National Primate Research Center (ONPRC) Tissue Distribution Program (TDP) at Oregon Health and Science University (OHSU) in Portland, Oregon. Three biological replicates (n=3) each from males and females were analyzed unless otherwise noted. Tissue homogenate preparation is described in detail in the Supporting Information.

Animal tissues were collected postmortem from pooled donors; samples were not donor matched across tissue types. NHP animals received ketamine (20 mg/kg) injection on average 15-30 min before necropsy; some animals may have also received isoflurane. NHP brain tissue samples were collected from prefrontal cortex and the primary visual cortex. NHP tissue samples were wrapped in foil and flash frozen or collected in Covaris tissueTUBE bags. Pathologists performed gross evaluation to establish that collected samples were from tissues within normal limits.

Only two biological replicates could be acquired for male human brain tissue; a pseudo-replicate was prepared by combining tissue from each of the two replicates to generate three total samples for statistical tests. One male human brain tissue sample was obtained from a drug overdose.

Whole tissue homogenate preparation

Animal and human tissues in ~1 g flash frozen portions stored at -80 °C were transferred to Covaris tissueTUBE TT2 bags. TT2 bags were maintained on dry ice until ready to pulverize. TT2 bags were chilled in liquid nitrogen for 5 seconds prior to pulverization on a Covaris CryoPREP CP02 at setting 4-5. Pulverized tissue was then transferred to a pre-weighed, pre-chilled 50 mL Falcon tube and weighed. For whole tissue homogenate, pulverized tissues were suspended in cold 250 mM sucrose in 1X PBS at a volume of 10 mL per 1 g liver or kidney tissue, or 5 mL per 1 g heart, lung, or brain tissue. The tissue suspension was homogenized using a mechanical tissue tearer over ice. Crude homogenates were centrifuged at 1,000 × g at 4 °C for 10 min to remove large debris, unbroken cells, and nuclei. Clarified homogenates were transferred via pipet to a fresh tube. Protein concentration was determined using a bicinchoninic acid (BCA) assay. Homogenates were aliquoted into single-use portions, flash frozen, and stored at -80 °C for future use.

Membrane fraction tissue homogenate preparation

Whole rat (Sprague Dawley) brains were transferred into pre-labeled, pre-weighed individual TT2 tissueTUBEs (Covaris, 520021) and tissue wet weights recorded. The tissue TUBESs were submerged in liquid nitrogen for 5-10 seconds immediately before inserting them one at the time into the cryoPREP automated dry pulverizer (Covaris, 500001). Each tissueTUBE was pulverized with using impact level 3. Submersion in liquid nitrogen and pulverization were repeated if the first impact did not pulverize the tissue into a fine powder. Once the brain was fully pulverized, the tissueTUBE was submerged in liquid nitrogen and transferred directly to a pre-chilled 15 mL dounce homgenizer tube. The brain was allowed to thaw on ice for 1-2 minutes. Once thawed, 10 mL of lysis buffer (filtered 20 mM HEPES pH 7.2, 1 mM DTT, 1 mM MgCl_2_, 2.5 U/mL benzonase) was added to the dounce homogenizer. Then, 15 passes were performed with the small pestle followed the large pestle, on ice. The resulting homogenate was transferred to 15 mL and centrifuged at 280 x g, 4 °C for 5 minutes to remove any residual large tissue debris. The supernatant was transferred to and ultracentrifuged for 45 minutes at 100,000 x g, 4 °C. The supernatant was removed, and the pellet was resuspended in 1 mL storage buffer (filtered 20 mM HEPES pH 7.2, 1 mM DTT) using a syringe equipped with a 21-gauge needle. The protein concentration of the resulting lysate was determined using Pierce^TM^ BCA protein assay (Thermo Fisher Scientific, 23227). The lysates were aliquoted into 1.7 mL tubes, flash-frozen in liquid nitrogen, and stored at -70 °C for subsequent probe labeling.

SDS-PAGE

Fentanyl analog solutions were handled in a chemical fume hood or ventilated class II biological safety cabinet (BSC). To generate samples for fluorescence SDS-PAGE analysis, click chemistry reagents were added in sequential order to 24 μL of the probe-labeled tissue homogenate: 0.5 μL of 1.5 mM 5-TAMRA picolyl azide (Click Chemistry Tools, 1254), 1 μL of 63 mM sodium-ascorbate, 1 μL of 25 mM THPTA ligand, and 1 μL of 50 mM copper (II) sulfate. Final click chemistry reagent concentrations were 30 μM, 2.5 mM, 1 mM, and 2 mM, respectively. After the addition of the click chemistry reagents, the samples were vortexed and placed in the thermal mixing blocks shaking at 500 rpm, 37 °C for 1 hour. After 1 hour incubation, 24 μL of 2x Tris-Glycine SDS sample buffer (Thermo Fisher Scientific, LC2676) and 5 μL of 10x NuPAGE^TM^ sample reducing agent (Thermo Fisher Scientific, NP0004) were added to the samples, followed by vortexing and incubation at 85 °C for 2 min. Samples (10 μL/well) were loaded into the protein gel (Invitrogen^TM^ Novex^TM^ WedgeGel^TM^   8-16 % Tris-Glycine 1 mm 15-well, Fisher Scientific, XP08165BOX) with 2 μL of Cytiva Amersham™ ECL™ Plex Fluorescent Rainbow Marker ladder (Cytiva RPN851E). Gels were run in 1X Tris-Glycine running buffer at 125 V for 2 hours before imaging with a Typhoon FLA 9500 laser scanner using the Cy3/Cy5 settings. After fluorescent imaging, the gels were fixed in 50% methanol, 7% acetic acid in Milli-Q water for 1 hour and stained with GelCode^TM^ Blue Stain Reagent (Thermo Fisher Scientific, 24590) for 1 hour while shaking. The gels were destained with 3 x 20-minute rinses in Milli-Q water before imaging on a Bio-Rad GelDoc EZ imager.

AfBPP sample preparation: click chemistry, enrichment, digestion, and TMT labeling

**Click chemistry:** To the remaining ~698 μL of the mass spectrometry sample, the following click chemistry reagents were added sequentially: 3 μL of 14 mM azido-PEG3-biotin conjugate in DMSO (Fisher Scientific, AAJ64996MC), 5 μL of 345 mM of freshly prepared sodium-ascorbate in water (Sigma, A4034-100G), 5 μL of 140 mM tris (3- hydroxypropyltriazolylmethyl) amine (THPTA) in water (Click Chemistry Tools, 1010), and 5 μL of 275 mM copper (II) sulfate in water (Alfa Aesar, A13986). Final click chemistry reagent concentrations were 60 μM, 2.5 mM, 1 mM, and 2 mM, respectively. After the addition of the click chemistry reagents, the samples were vortexed and incubated at 500 rpm, 37 °C for 1 hour. After incubation 700 μL of cold methanol was added to each sample and chilled in a -70 °C freezer for at least one hour. After being chilled for an hour, the samples were centrifuged at 14,000 x g, 4 °C for 15 minutes to pellet the precipitated proteins. The supernatant was removed/discarded, and the pellets were allowed to dry on the benchtop for 30 minutes. The pellets were then resuspended in 700 μL 1.2% SDS in 1X PBS via probe sonication with 18 x 1 second pulses (1 second sonication, 1 second pause) at an amplitude of 80%. Additionally, the samples were placed in the water sonication bath for 10 seconds. After the solutions were completely homogenized, the samples were heated at 95 °C for 2 minutes and then centrifuged at 14,000 x g for 5 minutes at 24 °C to pellet any insoluble material and protein concentrations were determined with Pierce^TM^ BCA protein assay (Thermo Fisher Scientific, 23227). The samples were stored at -70 °C until ready for enrichment.

**Enrichment:** The frozen samples were heated at 95 °C for 4 minutes, vortexed, and centrifuged at 14,000 x g for 5 minutes at room temperature to pellet any insoluble material. The clarified supernatants were then normalized with 1.2% SDS in 1X PBS to a final volume of 640 μL, to the lowest protein concentration amongst the sample for the tissue determined by the Pierce^TM^ BCA protein assay (Thermo Fisher Scientific, 23227) in 5 mL cryovials. All subsequent washes were performed using vacuum manifold (VWR, PAA7231). Chromatography columns of 1.2 mL bed volume (Bio-Spin ^®^, 7326025) were washed twice with 1 mL 1X PBS and 100 μL of streptavidin-agarose resin (Thermo Scientific, 20353) were added to the columns. The beads were washed 3x with 1 mL 0.5% SDS in 1X PBS, 3x with 1 mL 5 M urea in 25 mM HEPES (prepared fresh, pH 8), and 5x with 1 mL 1X PBS. After washing, beads were transferred twice per column with 1 mL of 1X PBS to the 4 mL cryovials containing the normalized proteome samples. After transferring the beads, 1.4 mL of 1X PBS was added to the cryovials so that we have roughly a 0.2% SDS concertation. The tubes were then rotated for 1 hour at 37 °C. After incubation, the protein-bound bead mixture was poured back into the columns. The cryovials were then rinsed with 1X PBS to transfer any remaining beads to the columns. This was repeated until there were no beads left in the vials. The protein bound beads were then washed 3x with 1 mL of 0.5% SDS in 1X PBS, 3x with 1 mL of 5 M urea in 25 mM HEPES (prepared fresh, pH = 8), 3x with 1 mL of MilliQ water, 9x with 1 mL 1X PBS, and 6x with 1 mL of 25 mM HEPES (pH 8). After washing, the protein-bound beads were transferred to 1.7 mL low binding tubes (Sorenson BioScience, 11500-E) with 2x 350 μL 5 M urea in 25 mM HEPES (prepared fresh, pH = 8). Then 28 μL of 100 mM TCEP-HCl were added to each tube and samples were reduced by incubating at 1,200 rpm, 37 °C for 30 minutes. Samples were then alkylated by adding 28 μL of 200 mM iodoacetamide and shaking at 1,200 rpm, 50 °C, for 45 minutes covered to shield from light. After reduction and alkylation, the protein-bound beads were transferred back to the corresponding column with 2x 500 μL volumes of 1X PBS. After transferring, the bound beads were washed 9x with 1 mL of 1X PBS and 5x with 1 mL of 25 mM HEPES (pH 8). After the final washing, the protein-bound beads were transferred to fresh 1.7 mL low binding tubes (Sorenson BioScience, 11500-E) with 2x 500 μL of 15 mM HEPES (pH 8). The samples were then centrifuged at 10,500 x g for 5 minutes at 24 °C and the supernatant was discarded.

**Digestion:** The protein-bound beads were resuspended in 210 μL of 15 mM HEPES (pH 8) and enough of 0.25 μg/μL trypsin (Thermo Scientific, 90057) was added to achieve a 1:4000 ration of μg trypsin to μg protein. Protein digestion was conducted overnight at 1,200 rpm and 37 °C. The following day, the samples were centrifuged at 10,500 x g for 5 minutes at room temp. After centrifugation, exactly 200 μL of the supernatant was transferred to low retention individually wrapped 1.5 mL tubes (Fisher Scientific, 02-681-331). Then 110 μL of HPLC grade water (Fisher Scientific, 14-650-357) was added to the beads and mixed at 1,200 rpm and 37 °C for 10 minutes. After mixing, the samples were centrifuged at 10,500 x g and 4 °C for 5 minutes and exactly 100 μL of the resulting supernatant was added to the corresponding samples in the individually wrapped tubes. A reference sample was prepared by combining 30 μL of the probe labeled replicates of each gender, then split into two tubes and 180 μL of HPLC grade water was added to each. Then 30 μL from the no-probe and competition samples was discarded to ensure similar drying times. The tubes were covered with Breathe-Easier membranes (Electron Microscopy Sciences, 7053720) and allowed to dry completely. The samples were then frozen in a -80 °C ultralow freezer.

**TMT labeling:** TMT labeling of samples was performed as previously described.^3^ Peptide quantity was determined using a fluorometric peptide assay (Fisher PI23290). The PQ samples were resuspended in 22 μL of 50% acetonitrile in HPLC grade water and incubated at 1200 rpm, 37 °C for 5 minutes. In a 96 well black round bottom costar plate 10 μL of the standards and samples were added. Then 70 μL of the Fluorometric Peptide Assay Buffer and 20 μL of the Fluorometric Peptide Assay Reagent were added to each well. The plate was then covered with a PCR film and incubated at room temperature for 5 minutes. The fluorescence intensity was measured at 390 nm and 475 nm using a BioTek Synergy plate reader, and the peptide concentration was calculated according to the manufacturer’s instructions.

The peptides were reconstituted in 20 μL of 50% acetonitrile in HPLC grade water. The pH was adjusted to 8.5 with 1 M HCl and incubated for 5 min at 1200 rpm and 37 °C. The TMT labeling reagents (Fisher PI90111 and PI90406) were reconstituted in anhydrous acetonitrile to a 17 μg/μL concentration solution. Then a volume of tag equal to 50x in 3 μL of the peptide concentration was added to each sample. The samples were then vortexed, spun down, and incubated at 25 °C for 1 hour at 400 rpm. Then the reactions were quenched by adding 2 μL of 5% hydroxylamine and incubated for 15 min at 25 °C and 400 rpm. After quenching, equal volumes of each sample in the same TMT-10 plex channel per tissue and sex were combined. The combined samples were evaporated to dryness in a SpeedVac concentrator. After the samples were dry, they were reconstituted in 100 μL of 8% formic acid/5% acetonitrile in HPLC water.

Stage tips were prepared by punching a 1 mm disk out of an Empore C18 extraction disk with a 16-gauge blunt needle and packing the plug into a 200 μL pipette tip. Each tip was placed in a 1.7 mL collection tube with a plastic washer. The stage tip was then conditioned by adding 100 μL MeOH and centrifugate at 1,000 rcf for 2 minutes at room temperature. Then the stage tips were washed with 100 μL of 80% acetonitrile/0.1% formic acid and centrifugated at 1000 rcf for 2 minutes at room temperature. After washing, the tips were equilibrated twice with 100 μL of 5% acetonitrile/0.1% formic acid and centrifugated at 1000 rcf for 2 minutes at room temperature. The pass through was discarded after each solvent addition and centrifugation. After the tips have been equilibrated, the combined samples were loaded into the tips and allowed to sit for 1 minute. Then the tips were centrifuged for 3 minutes at 1000 rcf and room temperature and pass through discarded. This was repeated if any liquid remained in the tips. The samples were then washed once with 100 μL 0.1% formic acid and twice with 100 μL of 5% acetonitrile/0.1% formic acid. After each wash addition the samples were centrifuged for 3 minutes at 1000 rcf and room temperature and the pass through discarded. The collection tubes where then exchanged for fresh 1.7 mL tubes for sample collection and 30 μL of the elution buffer (80% acetonitrile/0.1% formic acid), was added to the staging tips and let sit for 1 minute. The samples were centrifuged at 3000 rcf and room temperature for 1 minute. Then another 30 μL of elution buffer was added, allowed to sit for 1 minute, and centrifuged again at 3000 rcf and room temperature for 1 minute. The collected 60 μL was transferred to glass LC-MS vials and dried completely in a SpeedVac, and then resuspended in 12 μL of 3% acetonitrile/0.01% DDM in HPLC grade water.

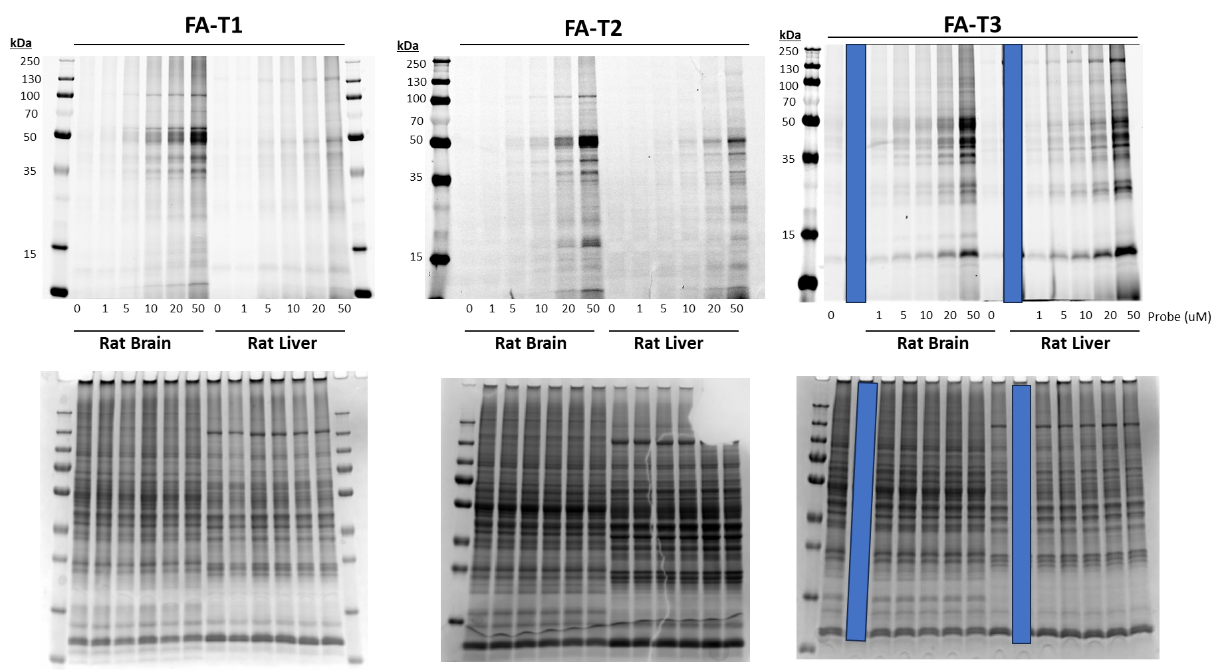

**Figure S4**. Dose dependent fentanyl AfBP labeling in rat brain and liver S9 fraction. Fluorescence (above) and GelCode Blue total protein stain (below) are shown for each gel.

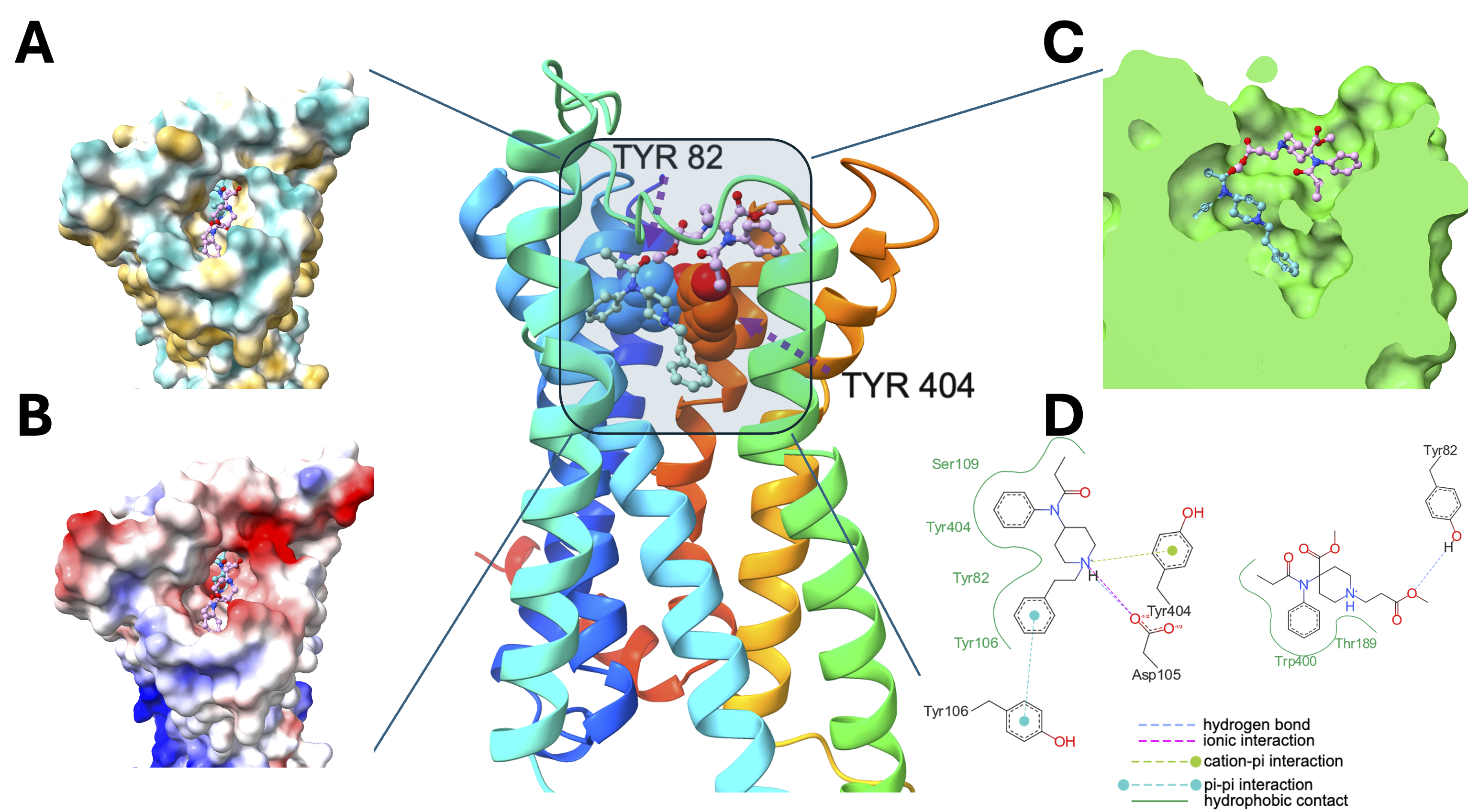

**Figure S5**. Docking simulation of fentanyl (light cyan colored ball and stick) and remifentanil (pink colored ball and stick) binding into muscarinic acetylcholine receptor M1 (CHRM1). Tyr82 is depicted as blue sphere, while Tyr404 is depicted as scarlet sphere. (A) Putative binding pocket in top view. Cyan: hydrophilic region, white: intermediate lipophilicity region, goldenrod: lipophilic region. (B) Same view as A, but with electrostatic potential visualization. Blue: positive, white: neutral, red: negative. (C) Ligands show feasible shape complementarity into binding pocket. (D) Fentanyl binding is driven by most common interactions (except metal-interaction) with nearby residues. Remifentanil binding is mainly driven by hydrophobic interactions with 1 hydrogen bond with Tyr82.
